# Role of Articulatory Motor Networks in Perceptual Categorization of Speech Signals: A 7 T fMRI Study

**DOI:** 10.1101/2023.07.02.547409

**Authors:** Kaisu Lankinen, Jyrki Ahveninen, Işıl Uluç, Mohammad Daneshzand, Azma Mareyam, John E. Kirsch, Jonathan R. Polimeni, Brian C. Healy, Qiyuan Tian, Sheraz Khan, Aapo Nummenmaa, Qing-mei Wang, Jordan R. Green, Teresa J. Kimberley, Shasha Li

**Author notes:** Corresponding author: Dr. Shasha Li, MD, Ph.D.: 149 13^th^ Street, Charlestown, MA, 02129, United States, Athinoula A. Martinos Center for Biomedical Imaging, Department of Radiology, Massachusetts General Hospital, Charlestown, MA, US, Harvard Medical School, Boston, MA, US.

## Abstract

**BACKGROUND:** The association between brain regions involved in speech production and those that play a role in speech perception is not yet fully understood. We compared speech production related brain activity with activations resulting from perceptual categorization of syllables using high field 7 Tesla functional magnetic resonance imaging (fMRI) at 1-mm isotropic voxel resolution, enabling high localization accuracy compared to previous studies.

**METHODS:** Blood oxygenation level dependent (BOLD) signals were obtained in 20 normal hearing subjects using a simultaneous multi-slice (SMS) 7T echo-planar imaging (EPI) acquisition with whole-head coverage and 1 mm isotropic resolution. In a *speech production localizer* task, subjects were asked to produce a silent lip-round vowel /u/ in response to the visual cue “U” or purse their lips when they saw the cue “P”. In a *phoneme discrimination task*, subjects were presented with pairs of syllables, which were equiprobably identical or different along an 8-step continuum between the prototypic /ba/ and /da/ sounds. After the presentation of each stimulus pair, the subjects were asked to indicate whether the two syllables they heard were identical or different by pressing one of two buttons. In a *phoneme classification task*, the subjects heard only one syllable and asked to indicate whether it was /ba/ or /da/.

**RESULTS:** Univariate fMRI analyses using a parametric modulation approach suggested that left motor, premotor, and frontal cortex BOLD activations correlate with phoneme category variability in the /ba/–/da/ discrimination task. In contrast, the variability related to acoustic features of the phonemes were the highest in the right primary auditory cortex. Our multivariate pattern analysis (MVPA) suggested that left precentral/inferior frontal cortex areas, which were associated with speech production according to the localizer task, play a role also in perceptual categorization of the syllables.

**CONCLUSIONS:** The results support the hypothesis that articulatory motor networks in the left hemisphere that are activated during speech production could also have a role in perceptual categorization of syllables. Importantly, high voxel-resolution combined with advanced coil technology allowed us to pinpoint the exact brain regions involved in both perception and production tasks.

## Introduction

The human brain has an astounding capability to extract meaning from acoustical signals and relate this sensory input to articulatory motor representations to produce speech. It is suggested that this process is facilitated predominantly by a left-hemispheric dorsal pathway (Hickok and Poeppel, 2007, Hickok and Poeppel, 2004). This pathway encompasses posterior parts of the superior temporal gyrus/sulcus (pSTG/pSTS), temporoparietal junction (TPJ), and articulatory motor areas of the frontal lobe (posterior inferior frontal regions, premotor cortex). According to this theory, during speech production, fast sensorimotor interactions across this “dorsal stream” help to validate the planned articulations by predicting the intended output based on the internal model of issued motor commands and actual sensory outcomes (Hickok et al., 2011). This circumvents the need for relying on monitoring of the auditory feedback from our own voice. However, whether the articulatory motor system is critically involved in speech perception remains unknown due to numerous incompatible findings (Hickok, 2009, Arsenault and Buchsbaum, 2015).

Over half a century ago, the motor theory of speech perception (MTSP) argued that production circuits are necessary for speech perception (Liberman et al., 1967). After falling from favor for many years, MTSP came back to the forefront after the discovery of “mirror neurons” in the primate motor cortices (Gallese et al., 1996). Consistent with MTSP, it was proposed that mirror representations of other speakers’ articulatory gestures could explain our ability to distinguish between phonemes, which have clear articulatory motor but ambiguous acoustic category boundaries (e.g., alveolar /da/ vs. labial /ba/). Evidence for categorical representations of speech in areas that control articulatory movements has since been obtained using neuroimaging and neuromodulation methods (Wilson et al., 2004, Pulvermuller et al., 2006, for a review, see e.g. Skipper et al., 2017). Transcranial magnetic stimulation (TMS) studies of the primary motor cortex (M1) suggest that the excitability of muscles controlling the tongue increases during listening to speech and during viewing speech-related movements (Fadiga et al., 2002, Watkins et al., 2003). TMS studies have also provided evidence for modulation of behavioral performance during speech sound processing, when stimulating articulatory-specific regions (D’Ausilio et al., 2009, Smalle et al., 2015, Mottonen et al., 2014a, Mottonen et al., 2014b, Schomers et al., 2015, Schmitz et al., 2019). Functional magnetic resonance imaging (fMRI) and magnetoencephalography (MEG) studies have in turn suggested that frontal speech production circuits are not only activated during speech perception tasks (Alho et al., 2016, Callan et al., 2010), but that the categorical attributes of speech sounds can be decoded using multivariate pattern analysis (MVPA) of signals from M1, premotor, and/or Broca’s areas (Schomers and Pulvermuller, 2016, Evans and Davis, 2015, Lee et al., 2012, Correia et al., 2015).

However, several critical arguments exist against the experimental evidence that supports MTSP. The perception-related effects in motor and premotor areas could be byproducts of the intrinsic connectivity between speech perception and production areas (Hickok, 2009), or they might reflect decision making rather than perceptual influences (Venezia et al., 2012). At the same time, the effects of TMS likely spread beyond the actual target areas both directly (i.e., point spread), and secondarily through long-range axonal connectivity. Some of the TMS findings, which were interpreted to support MTSP (Fadiga et al., 2002, Watkins et al., 2003, D’Ausilio et al., 2009, Smalle et al., 2015, Mottonen et al., 2014a, Mottonen et al., 2014b), could thus originate from beyond speech-related motor areas. The role of M1 could also be questioned based on the quite limited evidence of its direct efferent projections to auditory areas in mammals (as opposed to, e.g., premotor areas (Nelson et al., 2013, Schneider and Mooney, 2015). Furthermore, most fMRI studies published so far have been based on low anatomical resolution and/or exploratory rather than hypothesis-based decoding approaches. At conventional resolutions (2 mm or beyond), even individual voxels may capture signals across sulcal boundaries. This becomes particularly evident when using conventional volume-based approaches such as three-dimensional (3D) searchlights (Oosterhof et al., 2011) or univariate analyses using 3D smoothing kernels (Ahveninen et al., 2016a) (see **Figure 1**). Given these limitations, it thus remains unclear exactly which parts of articulatory motor areas have the most prominent association on speech-sound categorization.

**Figure 1.**
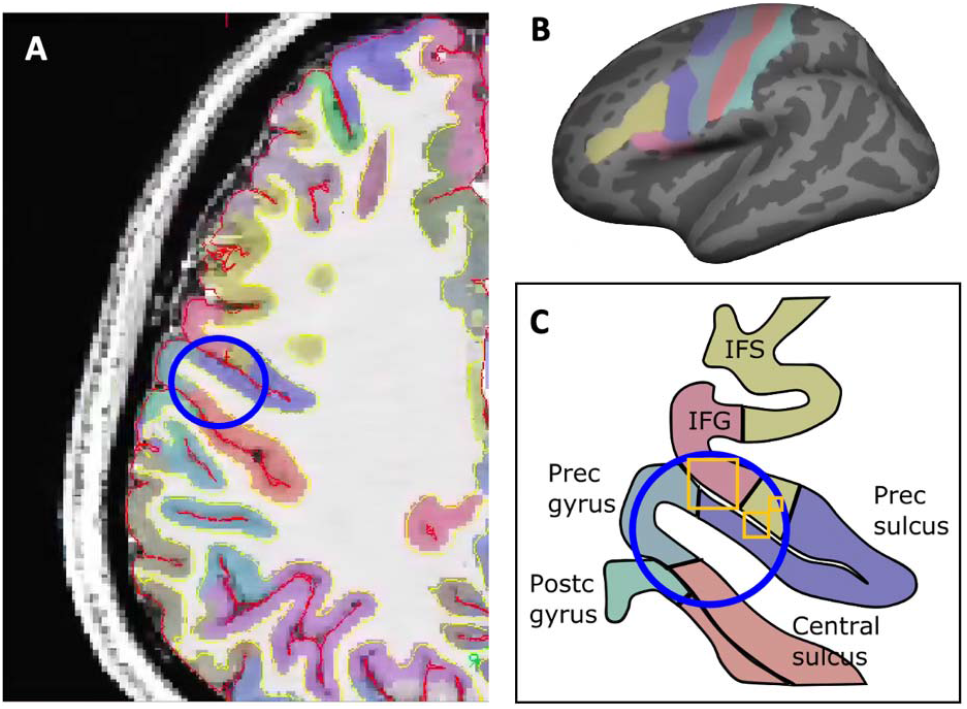
Effect of spatial resolution on the covered brain regions.**A**. An 8-mm (blue) radius sphere on the anatomical parcellation of one subject. This typically used radius intersects six different functional regions illustrated in **B**: inferior frontal sulcus (IFS), inferior frontal gyrus (IFG), precentral sulcus (Prec sulcus), precentral gyrus (Prec sulcus), central sulcus and postcentral gyrus (Postc gyrus). **C**. Schematic picture of the regions in **A** and **B** with an 8-mm (blue) radius sphere, and 3-mm, 2-mm and 1-mm voxels (orange squares).

Here, we aimed to pinpoint areas of articulatory motor pathways at high resolution using 7 T fMRI, which provides better sensitivity and signal-to-noise ratio than more conventional MRI field strengths (Edelstein et al., 1986, Yacoub et al., 2001, Ugurbil, 2018). Using a voxel size of 1-mm isotropic or below increases the specificity of fMRI signal within the gray matter, which helps prevent the spread of signals across adjacent sulci (Polimeni et al., 2010, Blazejewska et al., 2019, Ahveninen et al., 2016b). In addition, we used a custom-built 63-channel receive array coil, which overperforms the currently commercially available 32-channel coils (Mareyam et al., 2020). Combined high-resolution voxels and advanced coil technology introduces an advantage over previous works attempting to localize the processes involved in speech processing and perception. We used a combination of surface-based univariate general linear modeling (GLM) and region-of-interest (ROI) based MVPA (for a recent 7 T fMRI study with bigger voxel sizes, see (Archila-Melendez et al., 2018)) to localize the related regions. In the univariate analysis, we used a parametric modulation approach to find areas whose signal correlates with the phoneme category and the acoustic difference of the phonemes. In addition, we used MVPA to examine whether fMRI signals generated in areas associated with speech production can be utilized to classify perceived phoneme categories.

## Methods

### Subjects

The study protocol was approved by the Institutional Review Board at Massachusetts General Brigham (MGB), and a written or electronic informed consent form was obtained from all subjects prior to the experiments. Twenty healthy right-handed adults (12 females, 19–48 years, average 30 years) participated in the study. Potential subjects were screened to avoid MRI hazards and that they had ordinary hearing and (corrected) vision, and no developmental disorders, neurological diagnoses or medications influencing brain function. The participants were representative of the diverse local population, including a proportionate representation of minorities, as well as fluent in English language.

### Experimental Procedures

The subjects performed two types of tasks in the MRI scanner: two different sound **perception** tasks and a sound **production** task. For each of the tasks we collected two runs per subject. The length of each run was around 5 mins, and stimuli presented in the same order for each subject. For four subjects, we collected an additional run for the perceptions tasks.

In the sound **perception** tasks, we used stimuli modified from Smalle et al. (2015). We synthesized 350-ms loudness-matched syllables from an eight-step phonetic continuum between the phoneme prototypes /ba/ and /da/ using Praat (Boersma, 2001). This was accomplished by increasing the onset frequency of F2 from 1100 to 1615 Hz and that of F3 from 2250 to 2940 Hz, at eight equal base-2 logarithmic steps. Thus, we had phonemes /ba1/, /ba2/, /ba3/, /ba4/, /da1/, /da2/, /da3/ and /da4/, where /ba1/ and /da4/ were the most different, and /ba4/ and /da1/ were close to each other. All sounds were presented over MR-compatible, insert-style headphones (Sensimetrics Model S15, Malden, MA) at a comfortable volume level. To ensure that subjects understood the experimental procedures, each subject practiced the activation task prior to participating in the functional runs in the MRI scanner.

Each task consisted of trials whose stimulus order was balanced using optseq algorithm by FsFast to optimize the statistical fMRI deconvolution efficiency for our event-related design. The stimulus trials were interleaved with silent baseline trials (rest). We used two different **perception** tasks.

In the **speech production localizer** task, the subjects were asked to produce a silent lip-vowel sound /u/ in response to the visual cue “U” on the screen or to purse their lips in response to seeing the cue “P”. The aim was to use the same lip posturing with (/u/) and without (lip purse) association with speech sound production. There were 60 trials in one run, and the visual cues were presented randomly every 2.8–14 s.

In the **discrimination** task, the subjects listened to pairs of /ba/ and /da/ sounds (separated by stimulus onset asynchrony of 1 s), which were equiprobably either identical (e.g., /ba3/ vs, /ba3/ or/da1/ vs. /da1/) or separated by three intervals along the /ba/–/da/ continuum, e.g., /ba1/ vs. /ba4/ or /ba2/ vs. /ba5/. In one run, 69 stimulus pairs were (24 identical, 45 different) presented randomly every 2.8 s, except for 45 rest trials of length 2.8–11.2 s were randomly presented within one run. The subjects were asked to indicate whether they heard the same sound twice or two different sounds by pressing a button with their right-hand index (same sounds) and middle (different sounds) finger, respectively. They had time to respond until to the start of the next stimulus.

In **classification** task, subjects were presented with one of the eight stimuli of the /ba/–/da/ continuum at a time (number of trials within each condition balanced) and asked to indicate whether they heard /ba/ or /da/ by pressing a button with their right-hand index and middle finger, respectively. In one run, 79 stimulus trials were presented every 2.8–3.8 s. Occasionally, 17 rest trials of length 2.8–8.4 s were randomly presented within one run.

### MRI Data Acquisition

During scanning, the participants wore appropriate 7 T MRI compatible earphones for auditory stimuli and overlaid earmuffs to decrease the scanner noise. Head motion was minimized by firm support. The total duration of the imaging session was up to two hours.

7 T MRI data were obtained using a whole-body scanner (MAGNETOM Terra, Siemens, Erlangen, Germany). We used a custom-built 63-channel receive array coil (Mareyam et al., 2020) which uses a split helmet former contoured to the head on both sides and at the nape of the neck. The helmet former shown in **Supplementary Figure S1b** was sized to accommodate a majority of adult heads. The volume coil and top half of the receive array slide (in the bore direction) as shown in **Supplementary Figure S1c** to increase accessibility for the patient. **Supplementary Figure S1a** shows the receive array which has 24 elements on the top half (with a diameter of 5.5 cm) and 40 on the bottom half (with a diameter of 6 cm). The transmit coil is a detunable 16-rung band pass birdcage with a rung length of 24cm. The 64^th^ receiver channel is used for birdcage receive.

Anatomical T_1_ data were obtained using a 0.75-mm isotropic MEMPRAGE pulse sequence (van der Kouwe et al., 2008, Zaretskaya et al., 2018). fMRI was obtained by using a blipped-CAIPI simultaneous multi-slice (SMS) (Setsompop et al., 2012) echo-planar imaging (EPI) acquisition (gradient-echo T2*-weighted pulse sequence, TR/TE = 2800/27 ms, flip angle = 78°, fat suppression, FOV = 192 × 192 mm^2^, 192 × 192 matrix, 132 slices with thickness = 1.0 mm, bandwidth = 1446 Hz/pixel, acceleration factor in phase encoding direction 4, acceleration factor in slice encoding direction 3, nominal echo spacing 0.82 ms). The images were reconstructed online using the GRAPPA FLEET algorithm (Polimeni et al., 2016).

## Data Analysis

### Anatomical MRI preprocessing

Bias field correction, cortical surface reconstructions, coregistration of anatomical and functional data, and fMRI analyses were conducted using Freesurfer (Fischl, 2012) with an extension for submillimeter 7 T data (Zaretskaya et al., 2018) and our in-house software using Matlab 2019b. Surface reconstructions of the interfaces between the cortical gray matter vs. the underlying white matter and pial surface were automatically generated from the anatomical MRI data (Fischl, 2012).

### Functional MRI preprocessing

Distortions from the B_0_ field inhomogeneities were compensated by unwarping the fMRI volumes. Individual functional volumes were motion-corrected to the middle volume of each run, the fMRI volumes were coregistered with structural MRI using Boundary-Based Registration (Greve and Fischl, 2009), intensity normalized, and resampled into standard-brain cortical surfaces by projecting functional voxels onto the cortical surface vertices using trilinear interpolation. In addition, instead of a conventional volumetric smoothing approach, the resulting surface data were smoothed along the surface using a 2D Gaussian kernel with 3-mm, 6-mm and 10-mm FWHM. We used different degrees of smoothing to compare the effect of the parameter selection on the results and relate our small voxel-size analysis to the analyses using big voxels pooling activation from larger areas. This anatomically constrained smoothing has been shown to produce higher accuracy than volumetric smoothing (Blazejewska et al., 2019, Andrade et al., 2001, Jo et al., 2007, Kiebel et al., 2000). Slow trends in the data were removed by high-pass filtering at 0.003 Hz. The head motion was detected and corrected (within a run for univariate analysis, within a session for MVPA analysis) based on the automatic image registration algorithm.

Using 1-mm high-resolution functional imaging enables more accurate localization of brain activity compared with conventional protocols (Gardumi et al., 2016). **Figure 1** demonstrates the effect of spatial resolution on the covered ROIs. With 3-mm voxel resolution (the biggest of the three orange squares in panel C), it is possible that one voxel extends to several ROIs, whereas 1-mm resolution (the smallest orange square in panel C) enables focusing on a single region. Furthermore, using a conventional 8-mm radius (blue circle in panels A and C) in MVPA can include even more ROIs and prevent making inferences of the exact underlying functional areas.

### General linear model (GLM) analyses

In the sound **discrimination** task, we used a parametric modulation modeling approach in FsFast framework, as the categorical and acoustic information are partially overlapping. We formed regressors for categorical difference (1 for identical pair of syllables, 0 for different syllables), acoustic difference (−3 to 3, difference between /ba/ and /da/ in an 8-step continuum), offset and button press, with event durations 1.35 s, starting from the onset of the first stimulus of the pair. The categories of the regressors were defined based on the presented stimuli, not based on the subjects’ responses. The categorical and acoustic difference regressors were demeaned and were not highly correlated (r = 0.02). The calculated contrasts were offset, categorical difference, physical difference and button press.

In the speech **production** task, we used a simple GLM, with the regressors reflecting onsets and offsets of /u/ or lip purse trials. The calculated contrasts were /u/-sound vs. baseline, lip purse vs. baseline and /u/-sound vs lip purse (U-P).

For all the tasks, six motion parameters (roll, pitch, yaw, and displacement on the 3 axis) were included in the GLM analysis as nuisance regressors.

For both **discrimination** and **production** tasks, the effect size estimates of the relevant contrasts for each subject were submitted to a second-level GLM, with the statistical significance being confirmed using cluster-based simulation tests to avoid false positives (Greve and Fischl, 2018). The significance of the activation clusters was calculated with Monte Carlo simulations of 10,000 iterations, with cluster-forming threshold of *p* < 0.05, cluster-wise *p*-value 0.01, and positive effect for discrimination task, two-tailed effect for production task.

### MVPA analysis

For MVPA analysis, we combined the data from **classification** and **discrimination** tasks, which both included responses to each of the stimuli along the /ba/-/da/ continuum, but with different stimulus presentation timing patterns and motor response mappings (classification: index finger to /ba/, middle finger to /da/; discrimination: index finger for repeated /ba/ or /da/, middle finger for a /ba/-/da/ or /da/-/ba/ pair). The different stimulus timing for the tasks was taken into account in the regressors of the GLM model, which was calculated for concatenated tasks for each subject. The resulting contrast effect size values for /ba/ and /da/ categories were entered as input features to the MVPA.

The MVPA analyses were conducted in each subjects’ native volume space with no spatial smoothing. Functional regions-of-interest were defined by significance values of the group-level contrast for difference between /u/-sound and lip purse in the surface space, and then morphed to individual subjects’ volume space. There were 10 significant clusters in the left hemisphere and 9 in the right hemisphere. The significance level for cluster-wise correction for multiple comparisons was two-tailed *p* = 0.05.

MVPA was conducted using support vector machine (SVM) method implemented in libsvm (Chang and Lin, 2011) and provided in the COSMOMVPA package (http://www.cosmomvpa.org/) (Oosterhof et al., 2016) in MATLAB. Separately for each subject and 19 ROIs, an SVM classifier with a radial basis function kernel (selected based on (Song et al., 2011)) and cost equal to one (C = 1) was trained using data from (N_Runs_ − 1) runs and tested on a dataset from one run, employing four-fold cross-validation. In four subjects, from whom we had data from two additional fMRI runs available (one per task), the MVPA was based on six-fold cross-validation. Given the large number of small voxels in each ROI, we used principal component analysis (PCA) to reduce the number of features to principal components that explained 95% of the variance of the data (Song et al., 2011). In each cross-validation fold, training was conducted using a (N_Runs_ − 1) × 2 × N_PCs_ data set and tested on a 2 × N_PCs_ dataset.

To control for multiple comparisons, statistical significance of decoding accuracies in MVPA was tested using a nonparametric randomization approach. We used within-train-set cross-validation approach (Valente et al., 2021), where we created 500 random permutations so that the true labels of the classifier were shuffled within each exchangeability block (i.e., fold) in each training set. To determine the classification accuracies that emerge by chance with 2 classes, a null-distribution of decoding accuracies using training data with randomized item-content labels was calculated across all subjects and ROIs. For the final null distribution, we selected the maximum group mean decoding accuracy across all ROIs from each permutation. To assign a *p*-value for each connection, the original group mean accuracy value (found from classifiers with true labels) was compared with this null distribution.

## Results

### Behavioral analyses

The average discrimination accuracy (0.86 for same and 0.65 for different) in the **discrimination** task is shown in **Supplementary Figure S2** (left). Average responses across subjects and runs for the **classification** task are presented in **Supplementary Figure S2** (right). The fit of the average responses to a sigmoid function shows relatively sharp category boundary between categories /ba/ and /da/.

### GLM Analyses

Figure 2. shows the results from the univariate GLM analysis for the **discrimination** task. The activation related to categorical difference (differentiation between identical or different syllable pairs) was found in the left superior and inferior frontal cortex (especially pars opercularis), precentral sulcus and gyrus, superior postcentral regions and precuneus. Activation related inferior to the acoustic differences of the syllables was found in the right primary auditory cortex and to some extent in frontal areas. **Figure 2** also illustrates the methodological effect of smoothing parameters on the resulting activation clusters, maps becoming more focused when using smaller smoothing value.

**Figure 2.**
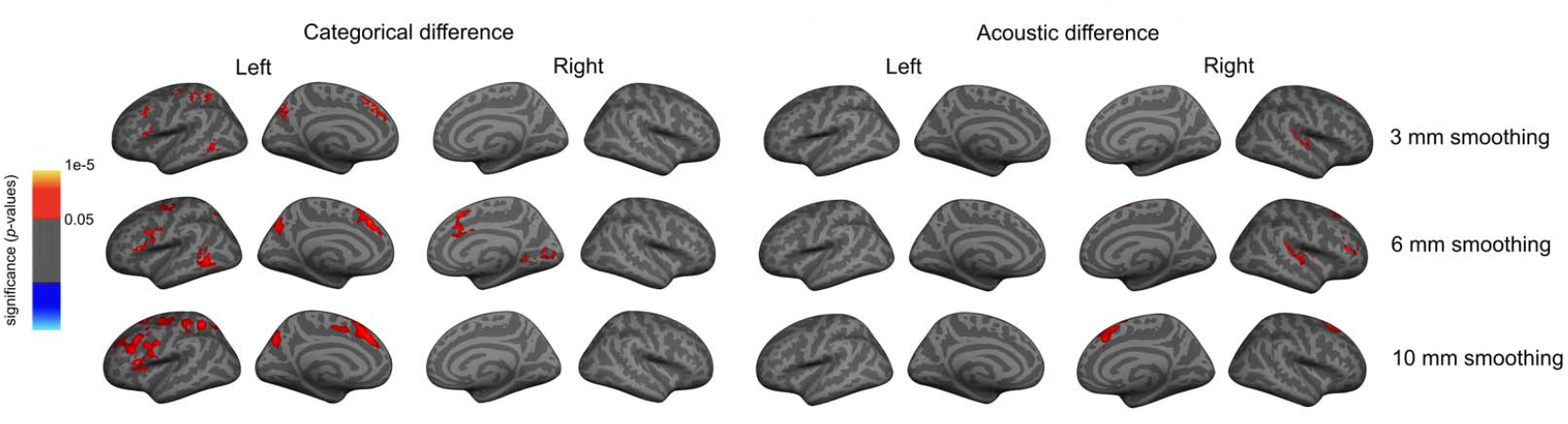
GLM results for categorical and acoustic difference regressors in the **discrimination** task with surface smoothing parameters 3 mm, 6 mm and 10 mm. The categorical difference reflects the differentiation between a pair of syllables being either identical or different (/ba/ or /da/). The acoustic difference is the distance from /ba/ to /da/ in an 8-step continuum of syllables. The color scale shows the significance values up to p = 1e-5. The significance of the activation clusters was calculated with Monte Carlo simulations of 10,000 iterations, with cluster-forming threshold of p < 0.05, cluster-wise p-value 0.01, and positive effect.

In the speech production task (**Figure 3**), the /u/-sound posture resulted in widespread activation bilaterally in inferior frontal, motor/premotor areas, and TPJ, whereas the lip purse resulted in less activation frontally and mostly in the right hemisphere. The difference between the /u/-sound lip posture and lip purse was most visible in the lower part of central sulcus bilaterally, and TPJ in the left hemisphere.

**Figure 3.**
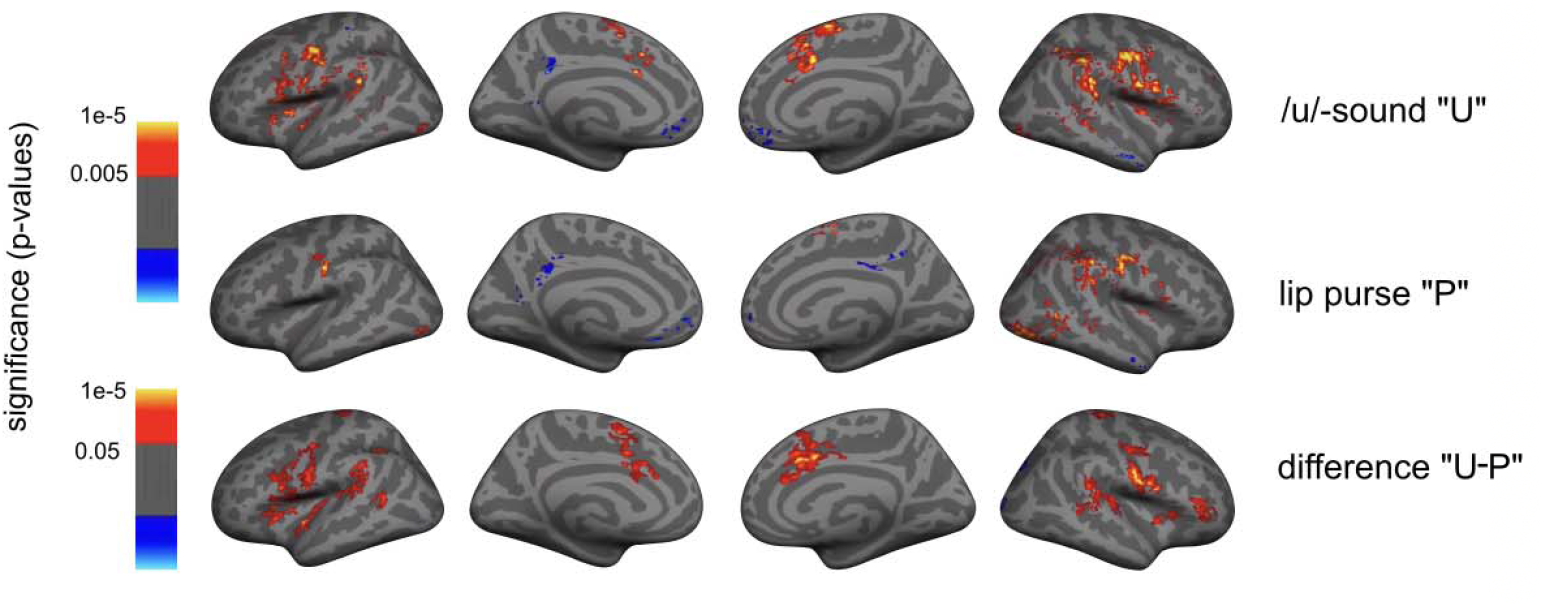
Speech production activity related to /u/-sound posture, lip purse and their difference with 3-mm surface smoothing (see 6 mm and 10 mm smoothing in **Supplementary Figure S3**). The significance of the activation clusters was calculated with Monte Carlo simulations of 10,000 iterations, with cluster-forming threshold of p < 0.05, cluster-wise p-value 0.01, two-tailed.

Our analysis shows that the activation patterns related to speech perception (**Figure 2**) and production (**Figure 3**) overlap in the inferior frontal regions and inferior precentral sulcus. This overlap suggests that the articulatory motor network has a role in speech perception, and that speech perception supports speech production.

### MVPA results

At the group level, the decoding accuracy was significantly above chance level (group mean 0.58, *p* < 0.022, maximum-statistic permutation test) in the cluster in the left precentral sulcus/pars opercularis (**Figure 4A, 4B**), which overlaps with motor and premotor regions controlling articulatory motor functions. The decoding accuracies of all 19 ROIs are shown in **Table 1. Figure 4C** shows the average confusion matrix for categorization of /ba/ and /da/ (see **Supplementary Figure S4** for individual subjects).

**Table 1.**
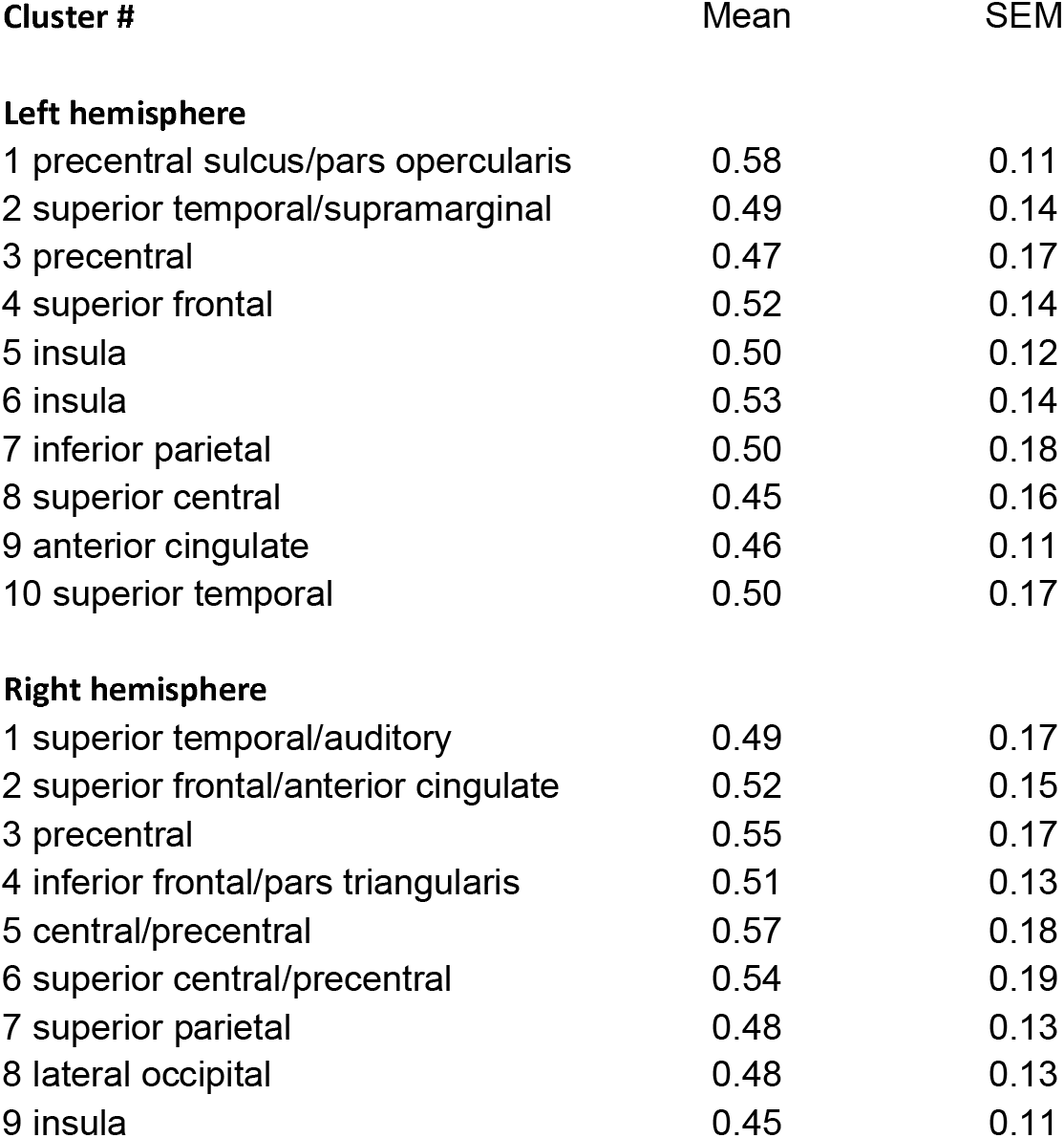
MVPA results. Group mean ± SEM (standard error of mean) decoding accuracies in all 19 ROIs (* *p* < 0.05 in a non-parametric permutation test).

**Figure 4.**
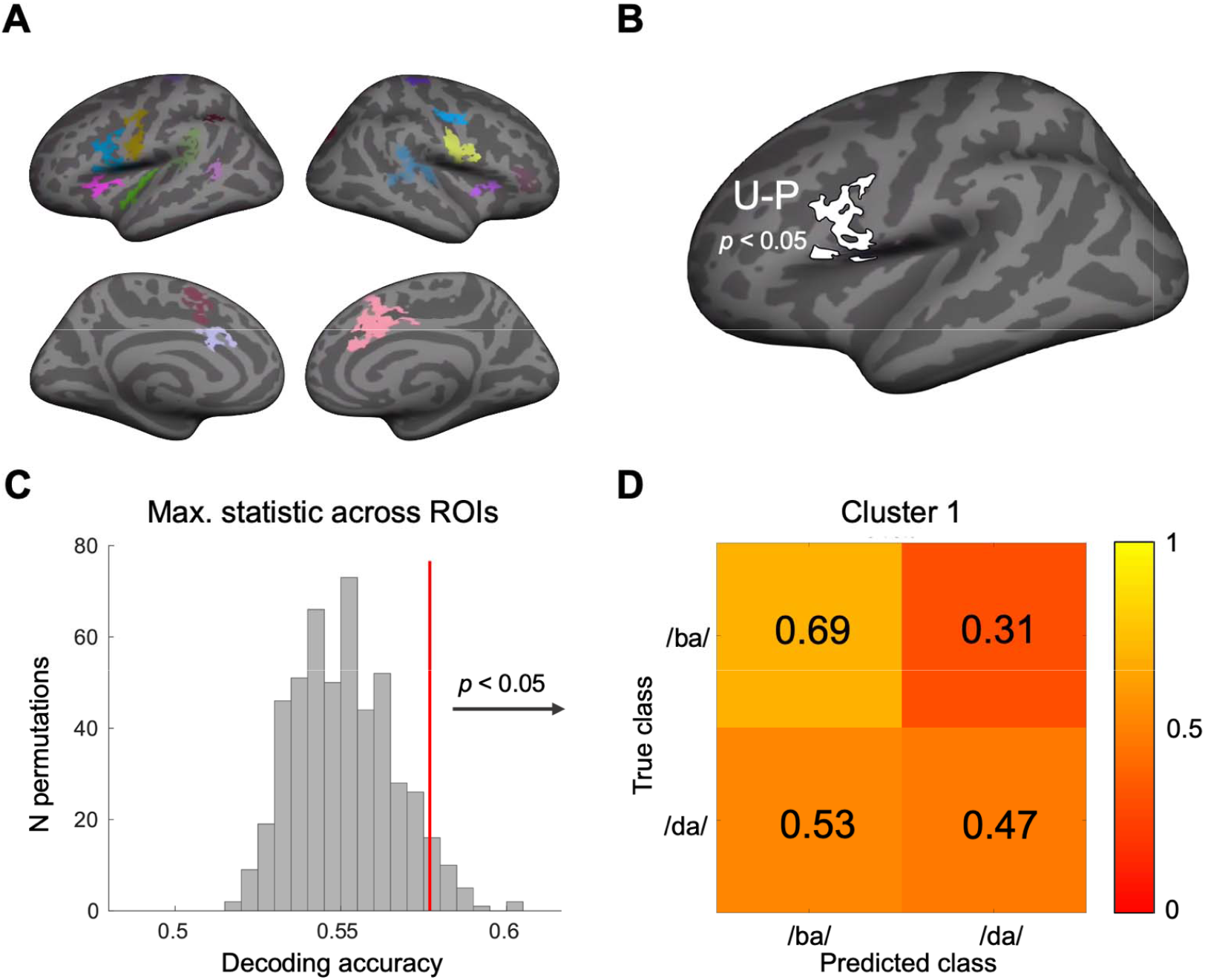
Summary of the MVPA results. A) The clusters based on the contrast U-P used in MVPA. B) The decoding accuracy was above chance level in cluster 1: the left precentral sulcus/pars opercularis region. C) Null distribution for 500 permutations. A vertical red line illustrates the critical value of 0.58 for p < 0.05. D) Confusion matrix for classification of /ba/ and /da/. Color scale indicates the accuracy of the classification (average across the subjects).

## Discussion

While it is established that the auditory system supports the articulatory motor network in speech production, the role of the articulatory motor areas in speech perception has been a highly debated subject (Hickok, 2010, Hickok et al., 2011, Pulvermuller and Fadiga, 2010, Scott et al., 2009, Skipper et al., 2017). Most of the evidence regarding the networks involved in speech production and perception are collected by anatomically typically inaccurate non-invasive neuroimaging methods, such as fMRI with large voxel-resolution, MEG, EEG, or TMS. We aimed at providing anatomically accurate information on the neural basis of the role of articulatory motor networks in speech perception using 1-mm resolution 7 T MRI acquisition. Our results demonstrated activation of the frontal and premotor areas during auditory syllable categorization tasks, which overlapped with the areas of speech production.

Our parametric-modulation univariate GLM analysis showed the relevance of left PreCS and PreCG in the syllable discrimination task. These regions correspond to the articulatory network in the “dorsal stream” model (Hickok et al., 2011). According to the model, the dorsal pathway is involved in processes related to sub-lexical segments of speech processing, such as syllables. The dorsal pathway is also recruited in phonological working memory, which is consistent with our discrimination task, where the comparison of two sequential sounds requires maintenance of the representation of the first syllable until the second one occurs. However, the “dorsal stream” model does not support the view that the motor networks would be necessary for speech perception, although it has been demonstrated that the precentral regions are activated during listening to speech sounds (Pulvermuller et al., 2006, Mottonen and Watkins, 2009, Mottonen et al., 2013). We found that the categorical difference activated premotor areas, slightly anterior to precentral sulcus. This difference from the previous studies could be explained by the lower resolution of the previous fMRI work, as well as the widespread activation by TMS, thus making the exact localization difficult.

Our MVPA analysis, which was conducted based on ROIs defined by the separate speech production localizer task, showed that the phonetic content can be decoded from the BOLD signals. Consistent with the univariate analysis, left precentral cortex/inferior frontal cortex was significantly associated with decoding accuracy using this approach. However, even though the decoding accuracy was the highest in this area, we cannot reject the possibility that other regions are included in the categorization process. However, in the present MVPAs, the classifiers were trained and tested across the two different tasks where the motor decisions (index finger vs. middle finger button press) were mapped orthogonally: In the classification task, the desired index finger response was associated with /ba/ and middle finger response with /da/ class. In the discrimination task, the index finger was the desired response for both /ba/-/ba/ and /da/-/da/ pairs and the middle finger for /ba/-/da/ and /da/-/ba/ pairs. Hence, it is very unlikely that the classification result is explainable by the contamination from a specific motor action.”

In the speech production localizer task, both the /u/-sound posture and lip purse activated central sulcus and inferior frontal regions. In general, producing a silent /u/-sound resulted in more widespread activation compared with lip pursing. Their difference was the most pronounced in the left precentral sulcus. One possible explanation for the widespread activation evident during the /u/-sound posture comes from studies that compare patterns of oromotor coordination during speech and non-speech behaviors (Moore, 1993). These studies have demonstrated that, for example, the upper and lower lip are rigidly coupled during lip pursing but more loosely coupled during /u/ (Ruark and Moore, 1997). The tight coupling of lip muscles activity during non-speech behaviors has been characterized as a less complex coordinative strategy than the diverse coupling patterns that are required to produce the wide variety of vocal tract shapes produced during speech (Green et al., 2000). More complex coordinative strategies, therefore, might recruit more neural resources. In addition, because /u/ is a vowel, it carries linguistic information, which might manifest in greater activation. The mutual activation of the inferior frontal regions in both speech perception and production suggests a role of the articulatory motor network in speech perception, although causality cannot be shown in the current experiment.

The use of simple sub-lexical syllables as stimuli limits the interpretation of our results. The brain processes involved in our tasks likely only partially overlap with those used in natural speech perception and language comprehension(Schomers et al., 2015, Schomers and Pulvermuller, 2016). Thus, future work with more naturalistic stimuli and experiments would be needed to obtain a more comprehensive picture of the processes involved in speech perception.

In conclusion, our findings were in line with previous studies demonstrating a relationship between articulatory motor networks and auditory speech perception. Importantly, high-resolution 7 T fMRI acquisition combined with advanced coil technology allowed us to investigate the cortical locations of the processes with high-spatial accuracy, enabling focusing on single regions-of-interest without significant spread to the neighboring areas. In future studies, this this high-resolution approach could be investigating sparse brain activity patterns, which have been suggested to constitute a fundamental organizational principle of human cognition (Jaaskelainen et al., 2022). At the same time, better understanding of speech perception and production could aid future research aiming at identifying precise biomarkers to aid in the prediction and management of communication deficits in various brain disorders.

## Acknowledgements

Our work was funded by the NIH grants K23DC018022, R01DC017991, R01DC016765, R01DC016915, and P41EB015896. We thank Dr. Lawrence L Wald, Dr. Lauryn Zipse, Grae Arabasz, Jacob Calkins, Alex Robertson, Dr. Marziye Eshghi, Brian Richburg, Tori Turpin and Jennifer Fiedler for their advice and support.

## References

Ahveninen, J., Chang, W. T., Huang, S., Keil, B., Kopco, N., Rossi, S., Bonmassar, G., Witzel, T. & Polimeni, J. R. 2016a. Intracortical depth analyses of frequency-sensitive regions of human auditory cortex using 7T fMRI. Neuroimage, 143, 116–127.

Ahveninen, J., Chang, W. T., Huang, S., Keil, B., Kopco, N., Rossi, S., Bonmassar, G., Witzel, T. & Polimeni, J. R. 2016b. Intracortical depth analyses of frequency-sensitive regions of human auditory cortex using 7TfMRI. Neuroimage, 143, 116–127.

Alho, J., Green, B. M., May, P. J., Sams, M., Tiitinen, H., Rauschecker, J. P. & Jaaskelainen, I. P. 2016. Early-latency categorical speech sound representations in the left inferior frontal gyrus. Neuroimage, 129, 214–23.

Andrade, A., Kherif, F., Mangin, J. F., Worsley, K. J., Paradis, A. L., Simon, O., Dehaene, S., Le Bihan, D. & Poline, J. B. 2001. Detection of fMRI activation using cortical surface mapping. Hum Brain Mapp, 12, 79–93.

Archila-Melendez, M. E., Valente, G., Correia, J. M., Rouhl, R. P. W., VAN Kranen-Mastenbroek, V. H. & Jansma, B. M. 2018. Sensorimotor Representation of Speech Perception. Cross-Decoding of Place of Articulation Features during Selective Attention to Syllables in 7T fMRI. eNeuro, 5.

Arsenault, J. S. & Buchsbaum, B. R. 2015. Distributed Neural Representations of Phonological Features during Speech Perception. J Neurosci, 35, 634–42.

Blazejewska, A. I., Fischl, B., Wald, L. L. & Polimeni, J. R. 2019. Intracortical smoothing of small-voxel fMRI data can provide increased detection power without spatial resolution losses compared to conventional large-voxel fMRI data. Neuroimage, 189, 601–614.

Boersma, P. 2001. Praat, a system for doing phonetics by computer. Glot International, 5, 341–345.

Callan, D., Callan, A., Gamez, M., Sato, M. A. & Kawato, M. 2010. Premotor cortex mediates perceptual performance. Neuroimage, 51, 844–58.

Chang, C.-C. & Lin, C.-J. 2011. LIBSVM: A library for support vector machines. ACM Trans. Intell. Syst. Technol., 2, 1–27.

Correia, J. M., Jansma, B. M. & Bonte, M. 2015. Decoding Articulatory Features from fMRI Responses in Dorsal Speech Regions. J Neurosci, 35, 15015–25.

D’Ausilio, A., Pulvermuller, F., Salmas, P., Bufalari, I., Begliomini, C. & Fadiga, L. 2009. The motor somatotopy of speech perception. Curr Biol, 19, 381–5.

Edelstein, W. A., Glover, G. H., Hardy, C. J. & Redington, R. W. 1986. The intrinsic signal-to-noise ratio in NMR imaging. Magn Reson Med, 3, 604–18.

Evans, S. & Davis, M. H. 2015. Hierarchical Organization of Auditory and Motor Representations in Speech Perception: Evidence from Searchlight Similarity Analysis. Cereb Cortex, 25, 4772–88.

Fadiga, L., Craighero, L., Buccino, G. & Rizzolatti, G. 2002. Speech listening specifically modulates the excitability of tongue muscles: a TMS study. Eur J Neurosci, 15, 399–402.

Fischl, B. 2012. FreeSurfer. Neuroimage, 62, 774–81.

Gallese, V., Fadiga, L., Fogassi, L. & Rizzolatti, G. 1996. Action recognition in the premotor cortex. Brain, 119 (Pt 2), 593–609.

Gardumi, A., Ivanov, D., Hausfeld, L., Valente, G., Formisano, E. & Uludag, K. 2016. The effect of spatial resolution on decoding accuracy in fMRI multivariate pattern analysis. Neuroimage, 132, 32–42.

Green, J. R., Moore, C. A., Higashikawa, M. & Steeve, R. W. 2000. The physiologic development of speech motor control: lip and jaw coordination. J Speech Lang Hear Res, 43, 239–55.

Greve, D. N. & Fischl, B. 2009. Accurate and robust brain image alignment using boundary-based registration. Neuroimage, 48, 63–72.

Greve, D. N. & Fischl, B. 2018. False positive rates in surface-based anatomical analysis. Neuroimage, 171, 6–14.

Hickok, G. 2009. Eight problems for the mirror neuron theory of action understanding in monkeys and humans. J Cogn Neurosci, 21, 1229–43.

Hickok, G. 2010. The role of mirror neurons in speech and language processing. Brain Lang, 112, 1–2.

Hickok, G., Houde, J. & Rong, F. 2011. Sensorimotor integration in speech processing: computational basis and neural organization. Neuron, 69, 407–22.

Hickok, G. & Poeppel, D. 2004. Dorsal and ventral streams: a framework for understanding aspects of the functional anatomy of language. Cognition, 92, 67–99.

Hickok, G. & Poeppel, D. 2007. The cortical organization of speech processing. Nat Rev Neurosci, 8, 393–402.

Jaaskelainen, I. P., Glerean, E., Klucharev, V., Shestakova, A. & Ahveninen, J. 2022. Do sparse brain activity patterns underlie human cognition? Neuroimage, 119633.

Jo, H. J., Lee, J. M., Kim, J. H., Shin, Y. W., Kim, I. Y., Kwon, J. S. & Kim, S. I. 2007. Spatial accuracy of fMRI activation influenced by volume- and surface-based spatial smoothing techniques. Neuroimage, 34, 550–64.

Kiebel, S. J., Goebel, R. & Friston, K. J. 2000. Anatomically informed basis functions. Neuroimage, 11, 656–67.

Lee, Y. S., Turkeltaub, P., Granger, R. & Raizada, R. D. 2012. Categorical speech processing in Broca’s area: an fMRI study using multivariate pattern-based analysis. J Neurosci, 32, 3942–8.

Liberman, A. M., Cooper, F. S., Shankweiler, D. P. & Studdert-Kennedy, M. 1967. Perception of the speech code. Psychol Rev, 74, 431–61.

Mareyam, A., Kirsch, J. E., Chang, Y., Madan, G. & Wald, L. L. A 64-Channel 7T array coil for accelerated brain MRI. Proceedings of the International Society for Magnetic Resonance in Medicine, 2020. 764.

Moore, C. A. 1993. Symmetry of mandibular muscle activity as an index of coordinative strategy. J Speech Hear Res, 36, 1145–57.

Mottonen, R., Dutton, R. & Watkins, K. E. 2013. Auditory-motor processing of speech sounds. Cereb Cortex, 23, 1190–7.

Mottonen, R., Rogers, J. & Watkins, K. E. 2014a. Stimulating the lip motor cortex with transcranial magnetic stimulation. J Vis Exp.

Mottonen, R., Van De Ven, G. M. & Watkins, K. E. 2014b. Attention fine-tunes auditory-motor processing of speech sounds. J Neurosci, 34, 4064–9.

Mottonen, R. & Watkins, K. E. 2009. Motor representations of articulators contribute to categorical perception of speech sounds. J Neurosci, 29, 9819–25.

Nelson, A., Schneider, D. M., Takatoh, J., Sakurai, K., Wang, F. & Mooney, R. 2013. A circuit for motor cortical modulation of auditory cortical activity. J Neurosci, 33, 14342–53.

Oosterhof, N. N., Connolly, A. C. & Haxby, J. V. 2016. CoSMoMVPA: Multi-Modal Multivariate Pattern Analysis of Neuroimaging Data in Matlab/GNU Octave. Front Neuroinform, 10, 27.

Oosterhof, N. N., Wiestler, T., Downing, P. E. & Diedrichsen, J. 2011. A comparison of volume-based and surface-based multi-voxel pattern analysis. Neuroimage, 56, 593–600.

Polimeni, J. R., Bhat, H., Witzel, T., Benner, T., Feiweier, T., Inati, S. J., Renvall, V., Heberlein, K. & Wald, L. L. 2016. Reducing sensitivity losses due to respiration and motion in accelerated echo planar imaging by reordering the autocalibration data acquisition. Magn Reson Med, 75, 665–79.

Polimeni, J. R., Fischl, B., Greve, D. N. & Wald, L. L. 2010. Laminar analysis of 7T BOLD using an imposed spatial activation pattern in human V1. Neuroimage, 52, 1334–46.

Pulvermuller, F. & Fadiga, L. 2010. Active perception: sensorimotor circuits as a cortical basis for language. Nat Rev Neurosci, 11, 351–60.

Pulvermuller, F., Huss, M., Kherif, F., MOSCOSO DEL PRADO Martin, F., Hauk, O. & Shtyrov, Y. 2006. Motor cortex maps articulatory features of speech sounds. Proc Natl Acad Sci U S A, 103, 7865–70.

Ruark, J. L. & Moore, C. A. 1997. Coordination of lip muscle activity by 2-year-old children during speech and nonspeech tasks. J Speech Lang Hear Res, 40, 1373–85.

Schmitz, J., Bartoli, E., Maffongelli, L., Fadiga, L., Sebastian-Galles, N. & D’Ausilio, A. 2019. Motor cortex compensates for lack of sensory and motor experience during auditory speech perception. Neuropsychologia, 128, 290–296.

Schneider, D. M. & Mooney, R. 2015. Motor-related signals in the auditory system for listening and learning. Curr Opin Neurobiol, 33, 78–84.

Schomers, M. R., Kirilina, E., Weigand, A., Bajbouj, M. & Pulvermuller, F. 2015. Causal Influence of Articulatory Motor Cortex on Comprehending Single Spoken Words: TMS Evidence. Cereb Cortex, 25, 3894–902.

Schomers, M. R. & Pulvermuller, F. 2016. Is the Sensorimotor Cortex Relevant for Speech Perception and Understanding? An Integrative Review. Front Hum Neurosci, 10, 435.

Scott, S. K., Mcgettigan, C. & Eisner, F. 2009. A little more conversation, a little less action--candidate roles for the motor cortex in speech perception. Nat Rev Neurosci, 10, 295–302.

Setsompop, K., Gagoski, B. A., Polimeni, J. R., Witzel, T., Wedeen, V. J. & Wald, L. L. 2012. Blipped-controlled aliasing in parallel imaging for simultaneous multislice echo planar imaging with reduced g-factor penalty. Magn Reson Med, 67, 1210–24.

Skipper, J. I., Devlin, J. T. & Lametti, D. R. 2017. The hearing ear is always found close to the speaking tongue: Review of the role of the motor system in speech perception. Brain Lang, 164, 77–105.

Smalle, E. H., Rogers, J. & Mottonen, R. 2015. Dissociating Contributions of the Motor Cortex to Speech Perception and Response Bias by Using Transcranial Magnetic Stimulation. Cereb Cortex, 25, 3690–8.

Song, S., Zhan, Z., Long, Z., Zhang, J. & Yao, L. 2011. Comparative Study of SVM Methods Combined with Voxel Selection for Object Category Classification on fMRI Data. PLOS ONE, 6, e17191.

Ugurbil, K. 2018. Imaging at ultrahigh magnetic fields: History, challenges, and solutions. Neuroimage, 168, 7–32.

Valente, G., Castellanos, A. L., Hausfeld, L., De Martino, F. & Formisano, E. 2021. Cross-validation and permutations in MVPA: Validity of permutation strategies and power of cross-validation schemes. Neuroimage, 238, 118145.

Van Der Kouwe, A. J., Benner, T., Salat, D. H. & Fischl, B. 2008. Brain morphometry with multiecho MPRAGE. Neuroimage, 40, 559–69.

Venezia, J. H., Saberi, K., Chubb, C. & Hickok, G. 2012. Response Bias Modulates the Speech Motor System during Syllable Discrimination. Front Psychol, 3, 157.

Watkins, K. E., Strafella, A. P. & Paus, T. 2003. Seeing and hearing speech excites the motor system involved in speech production. Neuropsychologia, 41, 989–94.

Wilson, S. M., Saygin, A. P., Sereno, M. I. & Iacoboni, M. 2004. Listening to speech activates motor areas involved in speech production. Nat Neurosci, 7, 701–2.

Yacoub, E., Shmuel, A., Pfeuffer, J., Van De Moortele, P. F., Adriany, G., Andersen, P., Vaughan, J. T., Merkle, H., Ugurbil, K. & Hu, X. 2001. Imaging brain function in humans at 7 Tesla. Magn Reson Med, 45, 588–94.

Zaretskaya, N., Fischl, B., Reuter, M., Renvall, V. & Polimeni, J. R. 2018. Advantages of cortical surface reconstruction using submillimeter 7 T MEMPRAGE. Neuroimage, 165, 11–26.

