## Supplementary figures S1-S4 for "Role of Articulatory Motor Networks in Perceptual Categorization of Speech Signals: A 7 T fMRI Study"

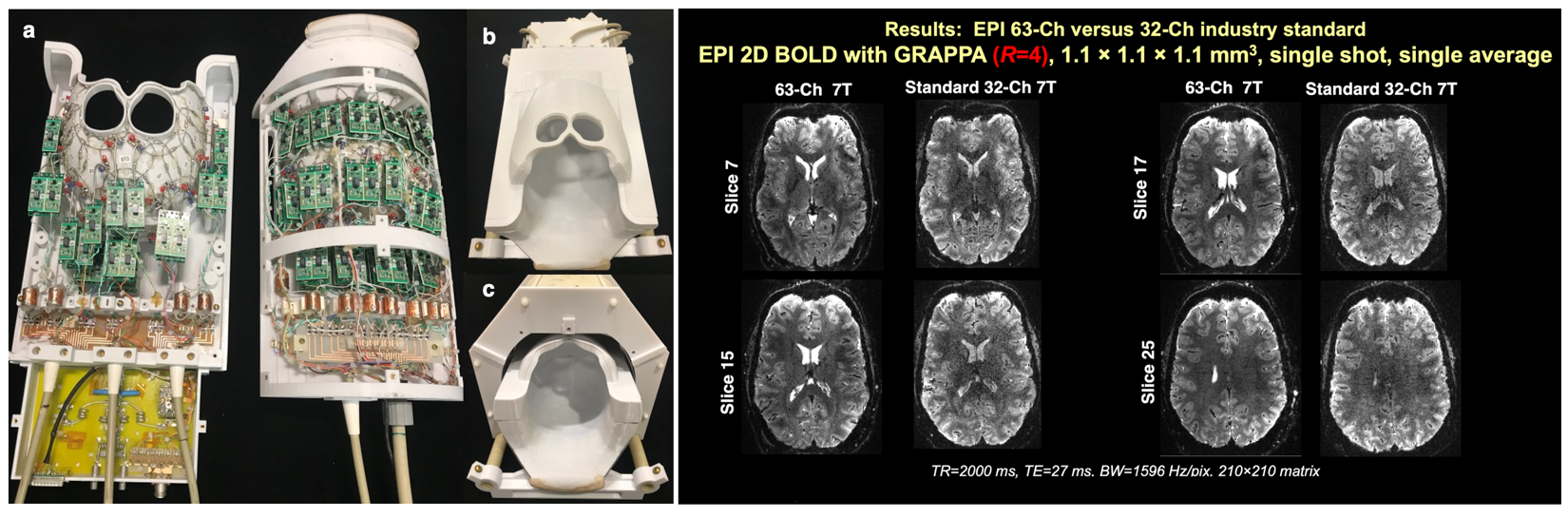

***Supplementary Figure S1.*** *(a) Elements laid out on formers with preamps mounted directly above coil detectors. (b) The two halves of the array with covers. (c) Array inside birdcage mounted on sliding rails.*

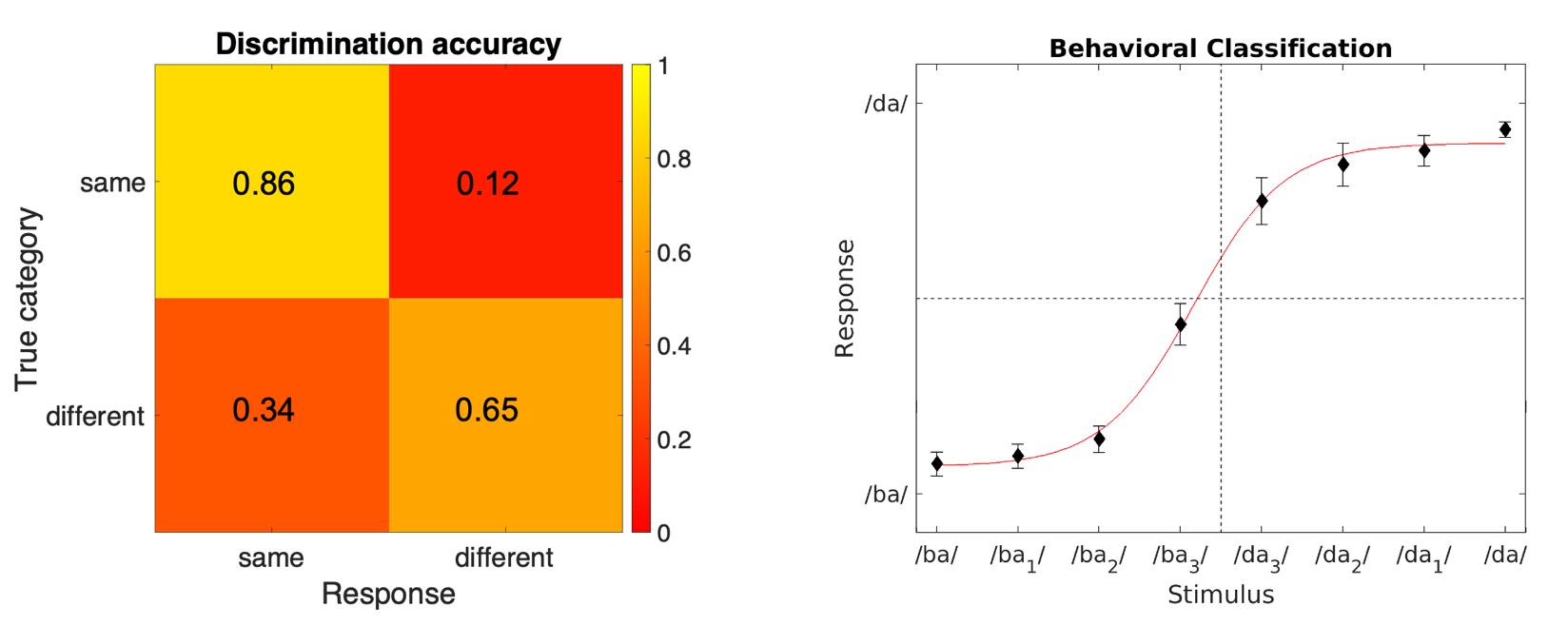

**a**

**b**

**c**

**Supplementary Figure S2**. **Left:** Discrimination accuracy for the **discrimination** task (average across subjects and runs). **Right:** Average responses across subjects and runs for the **classification** task (black diamond shape with standard error of means). The fit of the average responses to a sigmoid function shows relatively sharp category boundary between categories /ba/ and /da/.

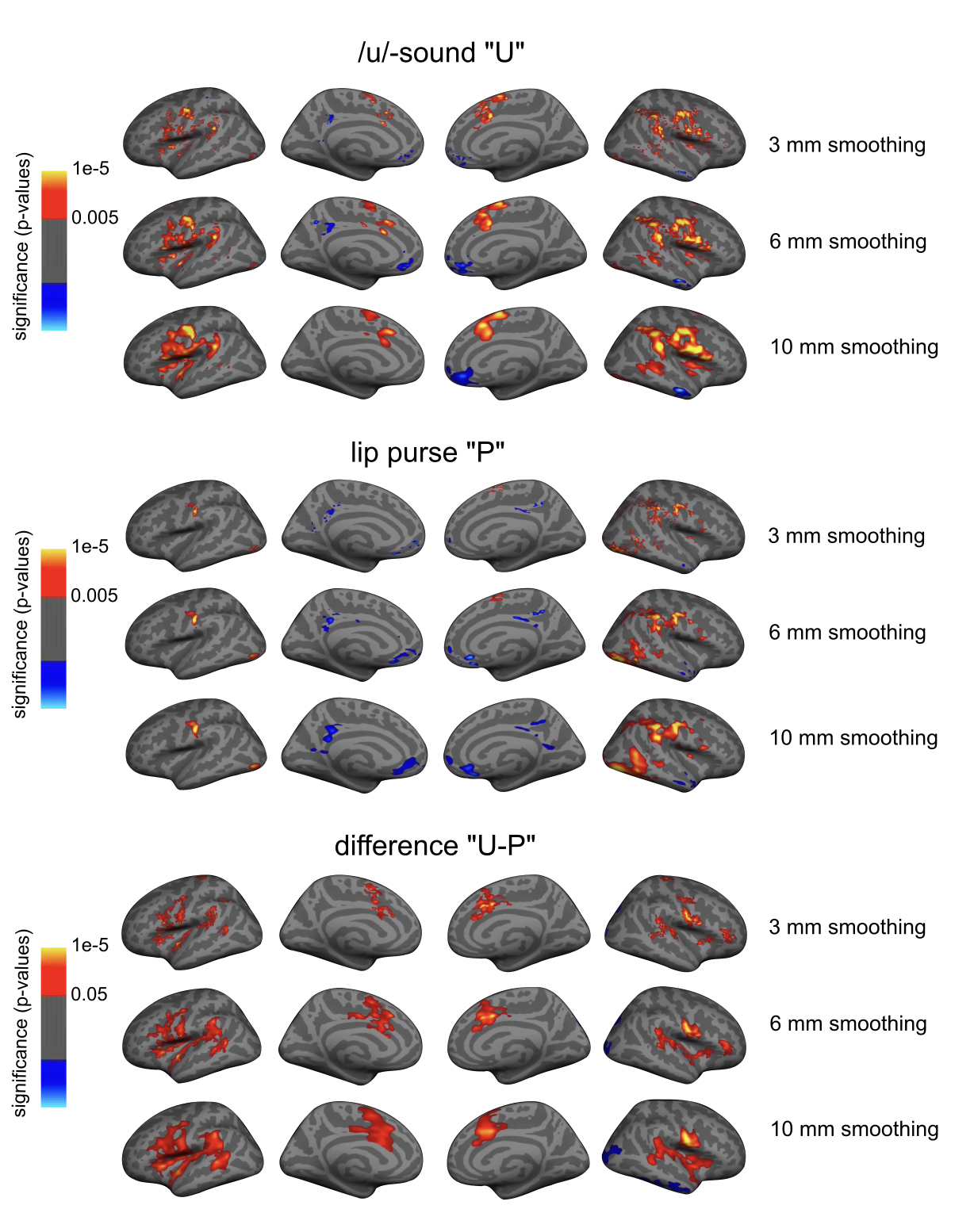

**Supplementary Figure S3.** Speech production activity related to /u/-sound posture, lip purse and their difference with 3 mm, 6 mm and 10 mm surface smoothing. The significance of the activation clusters was calculated with Monte Carlo simulations of 10,000 iterations, with cluster-forming threshold of *p* < 0.05, cluster-wise *p*-value 0.01, two-tailed.

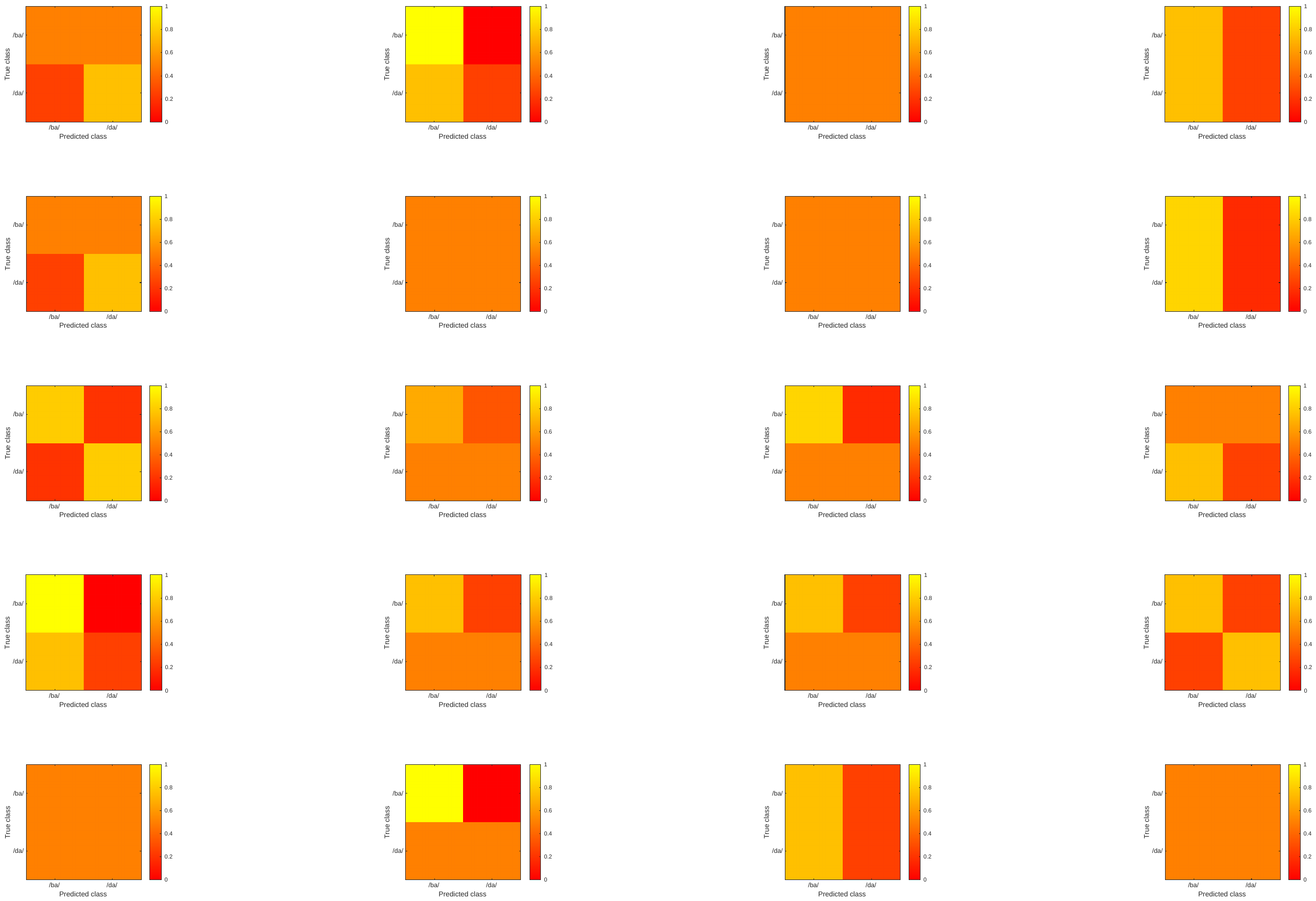

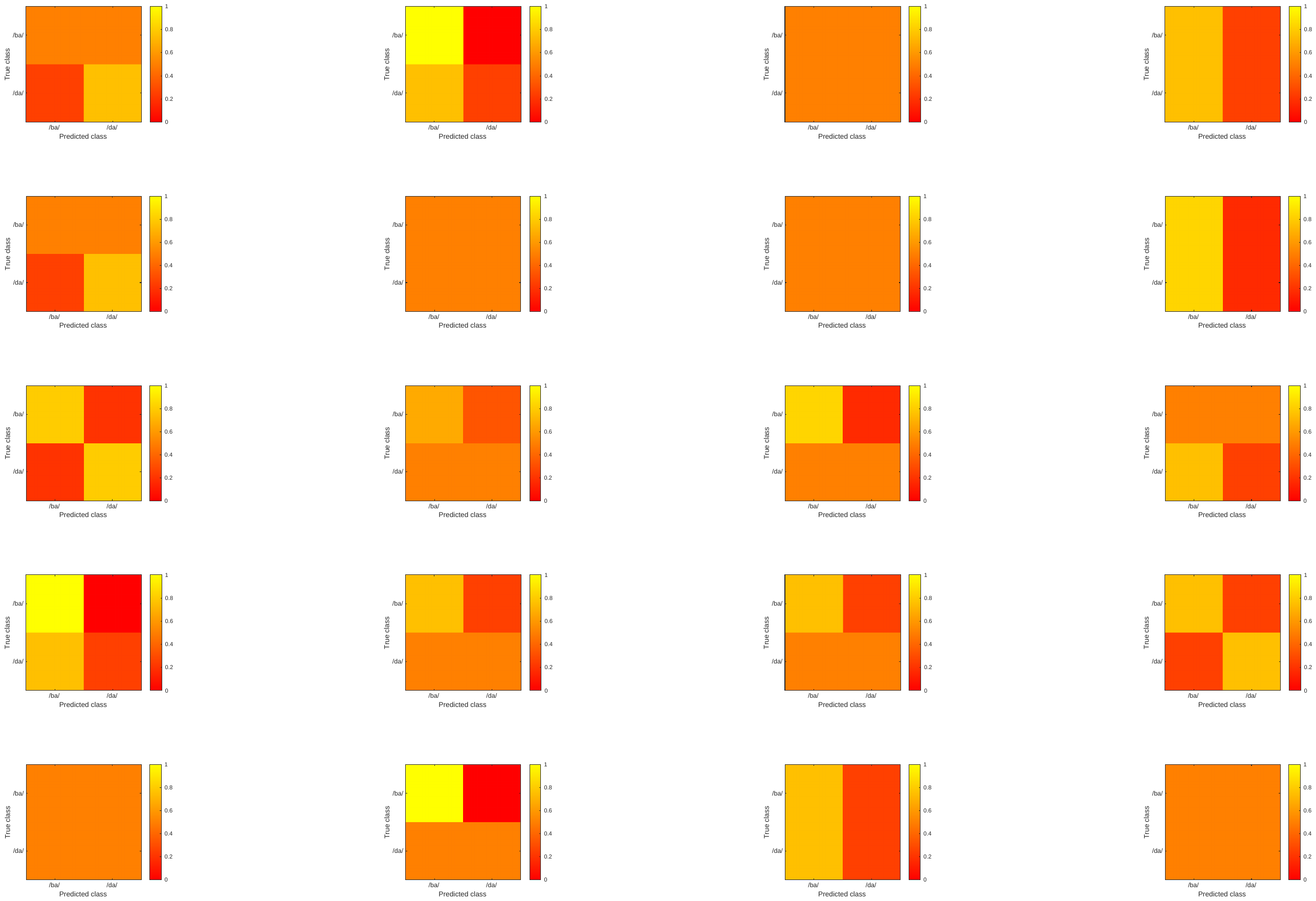

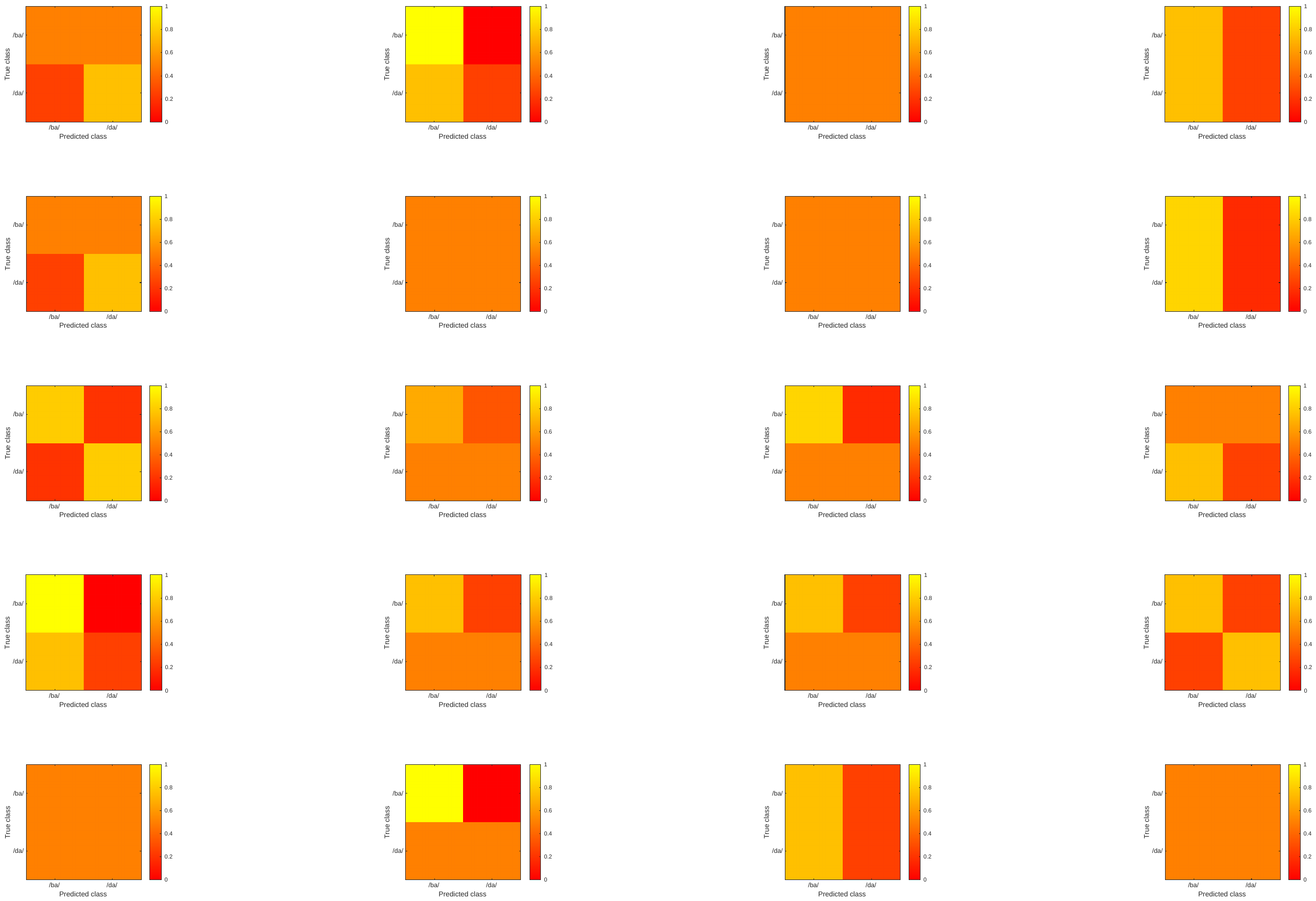

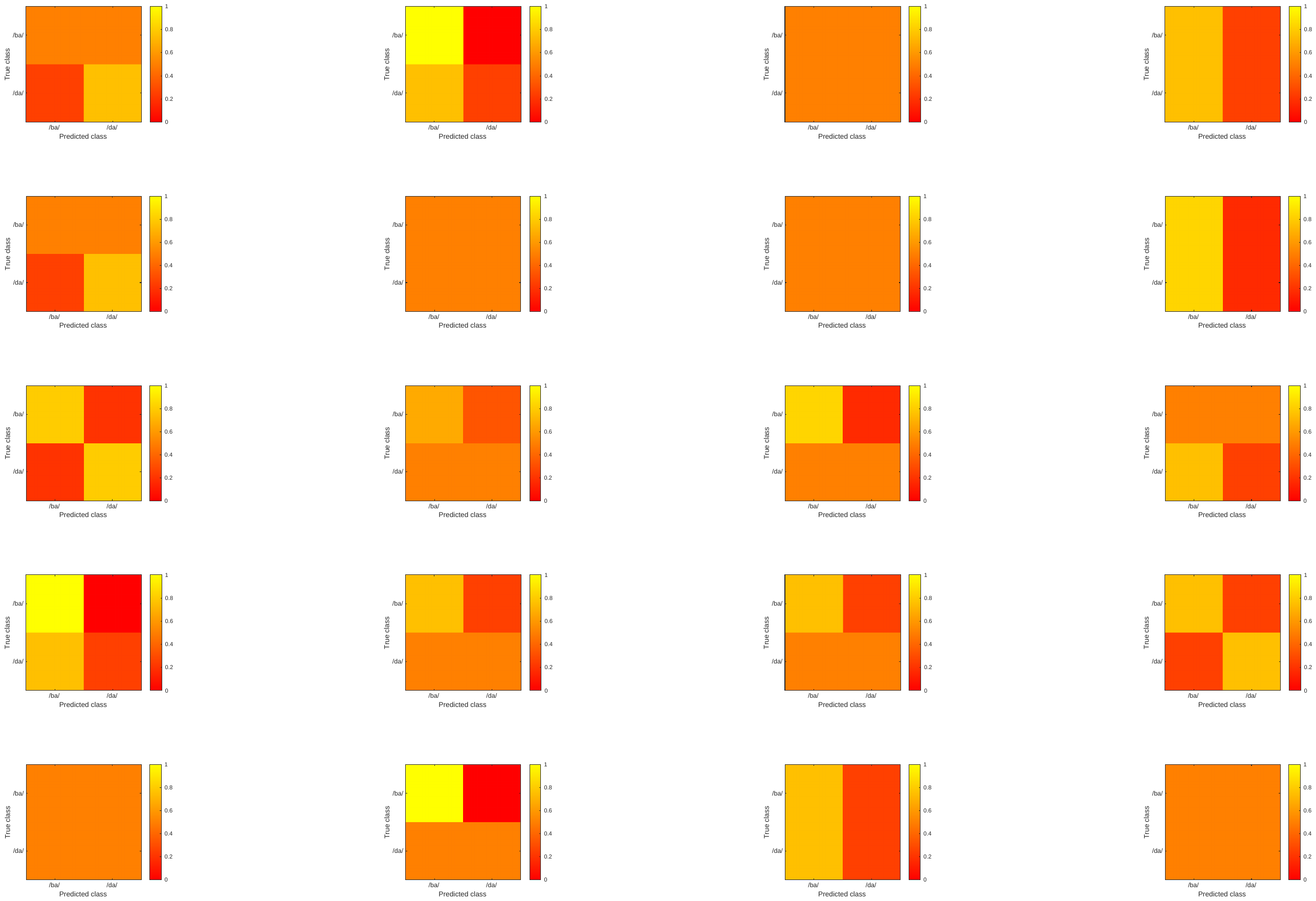

**Supplementary Figure S4**. Confusion matrices (predicted class vs. true class) for individual subjects from the MVPA categorization in the left inferior frontal cortex in cluster 1. Color scale indicated the accuracy of the classification.
